# Machine learning–based image analysis of Parkinson’s disease iPS cell-derived neurons predicts genotype and reveals mitochondria–lysosome abnormalities

**DOI:** 10.64898/2026.01.28.702423

**Authors:** Yan Li, Maximilian Powell, Jessica Chedid, Ratneswary Sutharsan, Adahir Labrador-Garrido, Dad Abu-Bonsrah, Chiara Pavan, Tyra Fraser, Dmitry Ovchinnikov, Melanie Zhong, Ryan Davis, Dario Strbenac, Jennifer A Johnston, Lachlan H Thompson, Deniz Kirik, Clare L Parish, Glenda M Halliday, Carolyn M Sue, Nicolas Dzamko, Gautam Wali

## Abstract

Mitochondrial and lysosomal dysfunction are central features of Parkinson’s disease (PD) across major genetic forms including *PRKN*, *SNCA*, and *LRRK2*. We applied cell morphomics, a machine-learning-based framework combining high-content imaging with quantitative feature extraction, to analyse mitochondrial and lysosomal morphology at single-cell resolution in iPS cell-derived cortical neurons from PD patients and healthy controls (13 lines total). Supervised machine-learning models distinguished PD neurons from controls with high accuracy (AUC = 0.87) and reliably separated individual genotypes. Feature importance and attribution analysis revealed genotype-specific organelle biases, with mitochondrial features dominating classification in *PRKN* neurons, balanced mitochondrial and lysosomal contributions in *SNCA* neurons, and a greater lysosomal contribution in *LRRK2* neurons. Multi-class models retained strong performance, and findings were reproduced across two independent laboratories using different dyes and imaging conditions. These results demonstrate that morphomics provides a robust and scalable framework to quantify genotype-specific organelle abnormalities in PD neurons and supports its application for cellular stratification and biomarker discovery.

## Introduction

Parkinson’s disease (PD) is clinically heterogeneous, with patients carrying different genotypes presenting variability in age at onset, rate of progression, and severity of motor and non-motor symptoms [1]. This heterogeneity is attributed to the downstream effects of genetic variation and their impact on cellular function.

Mitochondrial and lysosomal dysfunction are consistently observed across the most common genetic forms of PD - *PRKN*, *SNCA*, and *LRRK2*, with each mutation disrupting these cellular functions through distinct mechanisms[2]. *PRKN* mutations are classically associated with defective mitophagy and impaired clearance of damaged mitochondria, which secondarily compromises lysosomal function.

Dopaminergic neurons carrying *PRKN* mutations display reduced differentiation efficiency and neuritic complexity [3], impaired mitochondrial respiration and membrane potential [4–6], and defective mitophagy under oxidative stress conditions [7], together with enlarged lysosomes and disrupted mitochondria–lysosome contacts [8, 9]. *SNCA* mutations such as duplications and triplications are primarily associated with α-synuclein accumulation, that can initiate mitochondrial stress and imposes a secondary burden on lysosomal clearance. Dopaminergic neurons derived from *SNCA* lines show mitochondrial depolarisation, oxidative stress, and calcium dysregulation [10–12], followed by lysosomal enlargement, impaired trafficking, and reduced clearance of α-syn aggregates [11, 13, 14]. *LRRK2* mutations are linked to kinase-dependent defects in vesicular trafficking and lysosomal maturation, with secondary consequences on mitochondrial quality control and mitophagy. Neurons carrying *LRRK2* mutations consistently exhibit mitochondrial dysfunction and reduced membrane potential [15–17], together with impaired autophagosome transport and defective lysosomal processing driven by aberrant kinase activity [18, 19].

Given the consistent involvement of mitochondria and lysosomes across PD genotypes, we asked whether alterations in their morphology could be leveraged to classify and predict disease genotype and to determine if either organelle pathway is more prominently involved in each genotype. To achieve this, we used high-content confocal imaging coupled with computational analysis enabling the extraction of hundreds of quantitative features from thousands of single cells, generating multiparametric “morphomic” profiles that can be interrogated using machine learning to detect subtle multi-organelle phenotypes. Initial studies have provided proof-of-concept that cellular morphology encodes disease-relevant signatures. D’Sa *et al* profiled induced pluripotent stem (iPS) cell-derived cortical neurons and trained a machine-learning classifier on mitochondrial and lysosomal morphological features that distinguished PD subtypes including – chemically induced complex I dysfunction (rotenone-induced), induced mitophagy (oligomycin/antimycin-induced), proteinopathy (α-synuclein oligomer-induced), and patient-derived *SNCA* triplication and *PINK1* mutations [20]. Although this study demonstrated that morphomics can resolve both genetic and mechanistic subtypes, its conclusions were limited by using only a single line per group, restricting generalisability.

In this study, we sought to overcome these limitations by applying a morphomics framework to a multi-donor iPS cell-derived neuronal cohort spanning three major PD genotypes - *PRKN*, *SNCA*, and *LRRK2* with three independent patient lines per genotype and four healthy controls (13 lines in total). Using a Cell Painting–style approach [21, 22], we systematically profiled mitochondrial and lysosomal morphology at single-cell resolution. In total, 1,320 quantitative features per cell, capturing intensity, texture, and spatial organisation were extracted and used to train machine-learning classifiers to distinguish disease from control and between genotypes. To ensure robustness and generalisability, the workflow was validated independently across two laboratories using distinct mitochondrial stains and imaging conditions. This design enables not only accurate genotype classification but also quantitative attribution of whether mitochondrial or lysosomal pathways are more prominently affected in each genetic form of PD.

## Results

### Mitochondrial and lysosomal morphomic profiles distinguish PD from control cells

We first established a cell morphomics workflow to examine mitochondrial and lysosomal morphology at single-cell resolution (Figure 1). Patient- and control-iPS cell-derived neurons were stained to visualise mitochondria and lysosomes and imaged using high-content confocal microscopy. Quantitative features were extracted from these images (Supplementary figure 1) and used to train a machine-learning model to distinguish genetic PD samples from controls. A total of 1,320 quantitative features were extracted per cell (66,061 single cells analysed across the study). To understand which organelle variables drove the predictions, we applied SHAP (SHapley Additive exPlanations), which ranks features according to their influence on the model.

**Figure 1.**
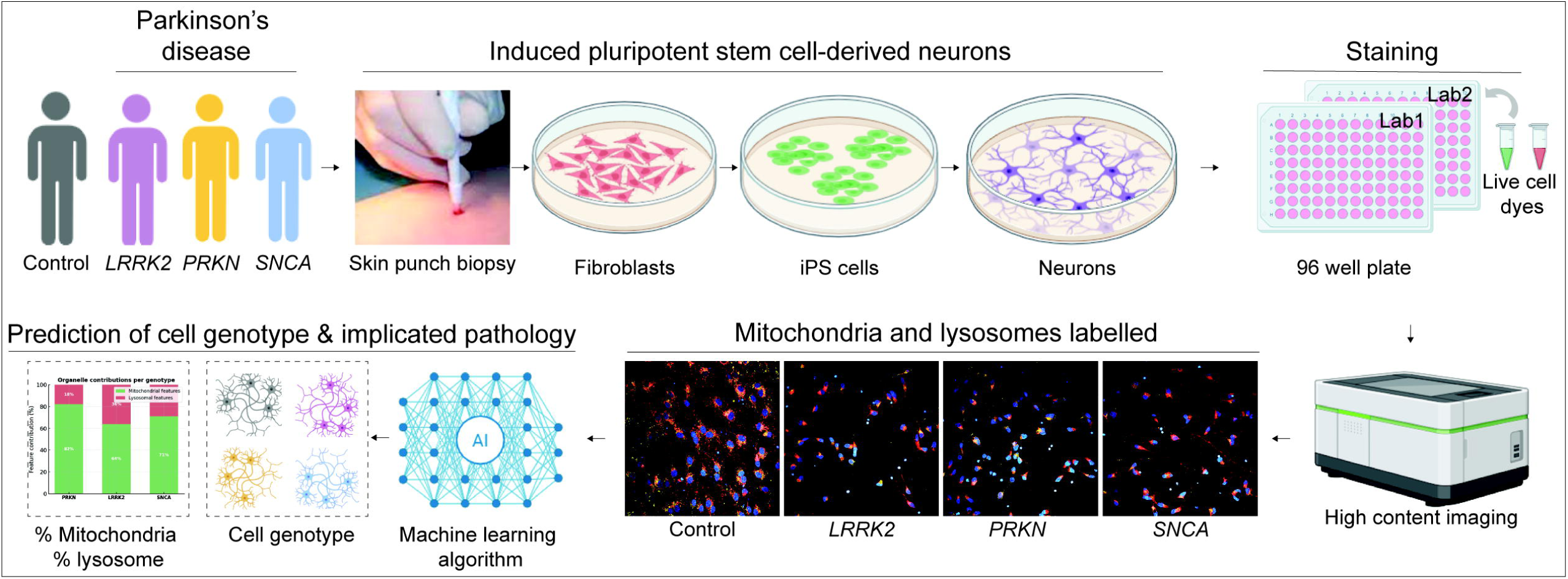
Cell morphomics workflow to identify Parkinson’s disease specific cellular signatures. iPS cell-derived neurons were seeded onto 96-well plates and labelled with live-cell dyes to visualise mitochondria and lysosomes. High-content imaging was performed to capture cellular morphology. Quantitative morphomic features were extracted and used to train an AI-based classifier that predicts cell genotype and reveals relative mitochondrial and lysosomal feature contribution underlying disease pathology.

The developed classifier achieved an AUC of 0.87 (95% CI: 0.87–0.88) (Figure 2A) with a balanced accuracy of ∼79% (82% of control cells and 75% of PD cells correctly identified) (Figure 2B). Performance was consistent across donors, indicating that the signal captures shared cellular changes. Misclassified cells may reflect overlap in morphology, with some control cells appearing stressed and some PD cells relatively preserved. SHAP-based feature attribution highlighted contributions from both mitochondria and lysosomes (Figure 2C). While the top 10 features were evenly divided, weighted SHAP contributions showed mitochondrial features account for a larger share of the model’s explanatory signal (∼59% vs 41% for lysosomal features) (Figure 2D). These proportions reflect SHAP-based feature attribution within the trained model and do not represent standalone classification performance of individual organelles. These results confirm that morphomic analysis can robustly distinguish PD from control cells and implicates both organelles in the disease phenotype.

**Figure 2.**
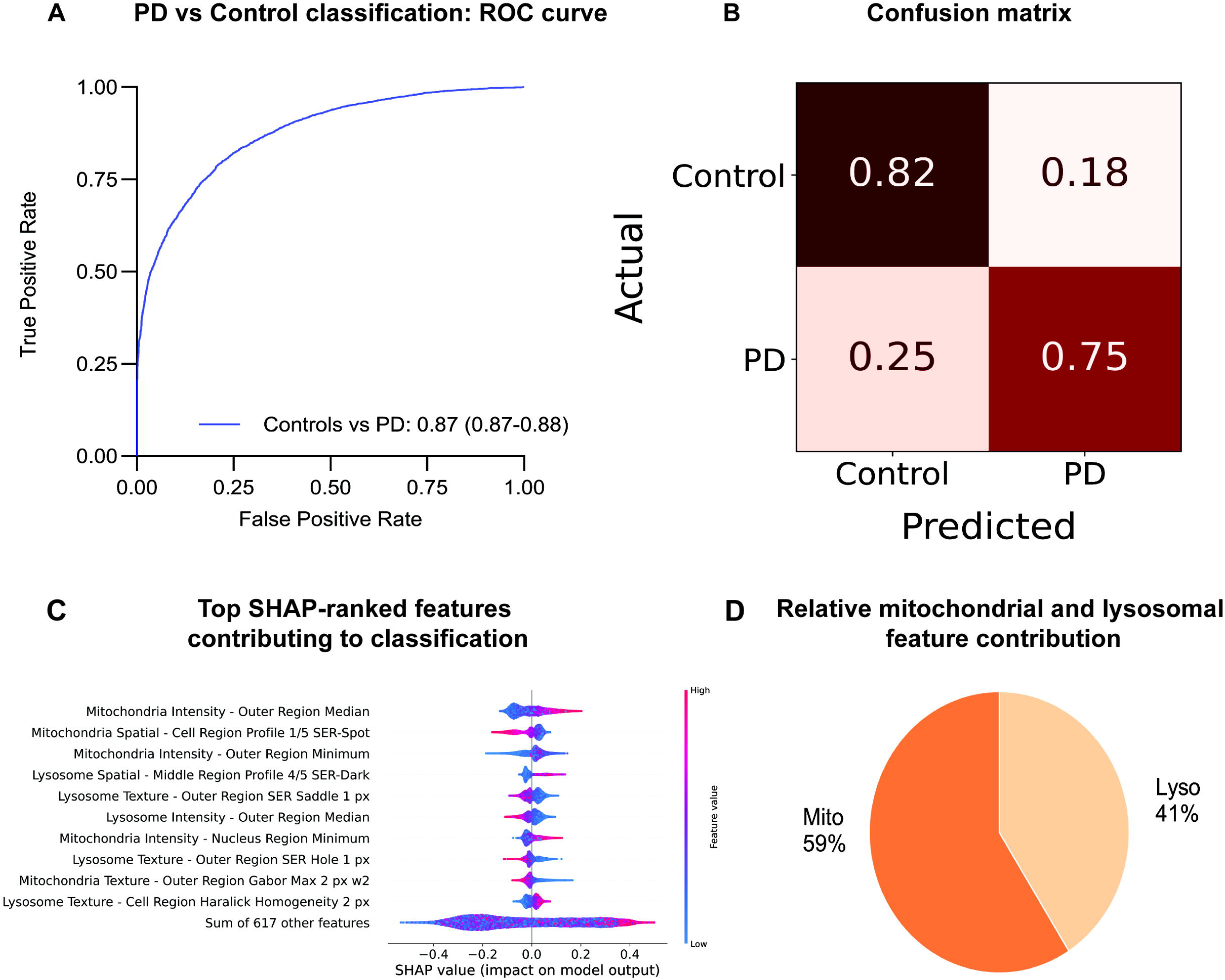
Classification of Parkinson’s disease neurons and relative organelle feature contribution. **(A)** ROC curve showing model performance in distinguishing Parkinson’s disease (PD) neurons from controls (AUC = 0.87, 95% CI 0.87–0.88) in the test dataset. **(B)** Confusion matrix illustrating model accuracy for PD and control classifications. **(C)** SHAP (SHapley Additive exPlanations) summary plot showing the top features contributing to model prediction, with mitochondrial and lysosomal intensity and texture parameters among the highest-ranked contributors. **(D)** Relative contribution of mitochondrial (59%) and lysosomal (41%) features to overall model performance, indicating that both organelle feature sets contribute substantially to disease classification.

To further interpret the organelle-level features driving classification, we examined the SHAP summary plot (Figure 2C). In this plot, features positioned toward the right (positive SHAP values) increase the model’s predicted probability of PD, whereas features positioned toward the left (negative SHAP values) increase the predicted probability of control; point colour represents the relative feature value, ranging from low (blue) to high (red). The morphological features using the “cell-painting” style approach capture patterns of staining intensity, spatial distribution, and texture (i.e., differences in pixel intensity distribution), enabling quantitative description of cellular phenotypes without explicit organelle segmentation.

Analysis of the top SHAP-ranked features indicates that mitochondrial and lysosomal contributions to classification arise from distinct morphomic signatures, with features from the outer cellular region (peripheral cell region, furthest away from nucleus) contributing prominently for both organelles. Highly ranked mitochondrial features were predominantly intensity-based (e.g. Outer Region Minimum and Median), indicating alterations in how mitochondrial signal is distributed across the cell. In contrast, highly ranked lysosomal features were largely texture-based (e.g. SER and Haralick descriptors), again primarily derived from the outer cellular region, reflecting differences in the spatial organisation and pixel intensity distribution of lysosomal staining.

### Genotype-specific separation and organelle attribution

The morphomics workflow was also able to distinguish each major genetic form of PD from controls. Classifier performance was strong across all comparisons, with AUC = 0.88 (95% CI: 0.87-0.89) for *PRKN* vs Control, 0.94 (95% CI: 0.93-0.95) for *LRRK2* vs Control, and 0.90 (95% CI: 0.9-0.91) for *SNCA* vs Control (Figure 3A). Accuracy was balanced in each case, with roughly 76–88% of cells correctly classified, showing that the cellular features captured by morphomics are sufficiently distinctive to resolve genotype-specific signatures (Figure 3B).

**Figure 3.**
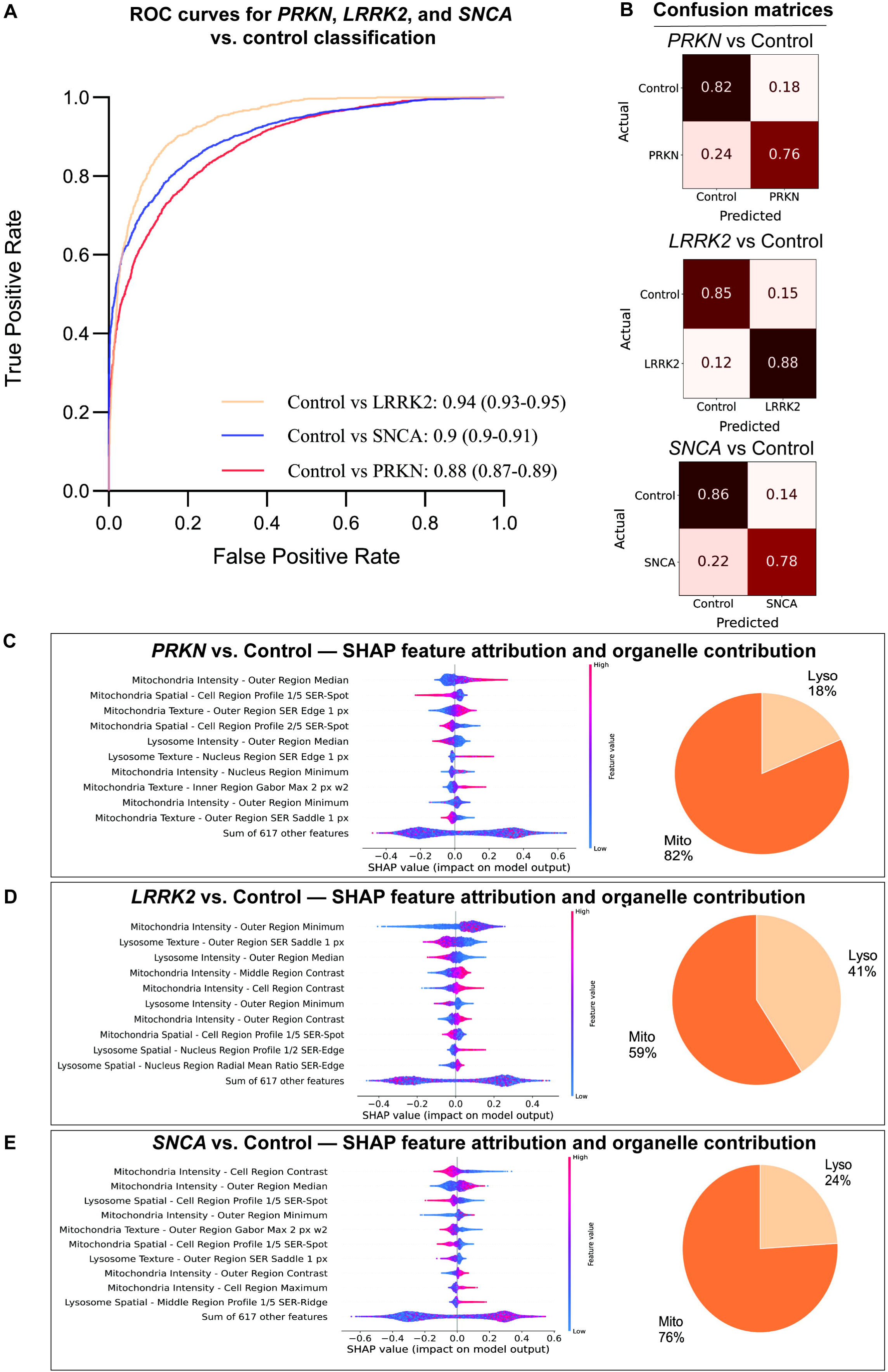
Genotype-specific classification and organelle feature attribution in Parkinson’s disease. **(A)** ROC curves showing machine-learning model performance in distinguishing *PRKN*, *LRRK2*, and *SNCA* patient-derived neurons from controls in the test dataset. Classification accuracy was highest for *LRRK2* (AUC = 0.94, 95% CI 0.93–0.95), followed by *SNCA* (AUC = 0.90, 95% CI 0.90–0.91) and *PRKN* (AUC = 0.88, 95% CI 0.87–0.89). **(B)** Confusion matrices illustrating prediction accuracy for each genotype relative to controls. **(C–E)** SHAP feature attribution plots and organelle contribution charts for *PRKN* (C), *LRRK2* (D), and *SNCA* (E) neurons. Mitochondrial features were the dominant contributors to model prediction, accounting for 82%, 59%, and 76% of explanatory power in *PRKN*, *LRRK2*, and *SNCA* neurons, respectively, with lysosomal features contributing the remainder.

Feature attribution analysis revealed organelle-specific differences across genotypes (Figure 3C–E). Weighted SHAP contributions showed that mitochondrial features accounted for the largest share in *PRKN* (82%), were moderate in *SNCA* (76%), and lowest in *LRRK2* (59%). Lysosomal features followed the opposite pattern, contributing 18% in *PRKN*, 24% in *SNCA*, and 41% in *LRRK2*. Across all genotypes, the most informative features for both organelles were predominantly derived from the outer cellular region, defined as the cell area distal to the nucleus (Figure 3C-E). *PRKN* cells showed a predominantly mitochondrial-associated signature, with highly ranked features reflecting changes in mitochondrial staining intensity in the outer cellular region, indicating altered mitochondrial distribution in the cell periphery. *LRRK2* cells, in contrast, showed a relatively stronger lysosomal-associated signature, with top features enriched for lysosomal texture and intensity measures from the outer cellular region. *SNCA* cells displayed an intermediate profile, with both mitochondrial intensity-based features and lysosomal texture-based features from the outer cellular region contributing to classification, indicating concurrent alterations affecting both organelles.

### Multi-modal classification across PD genotypes

We next evaluated a multi-class model to discriminate between *PRKN*, *LRRK2*, and *SNCA* genotypes. The classifier achieved AUCs of 0.87 (95% CI: 0.86-0.88) for *PRKN vs LRRK2*, 0.89 (95% CI: 0.88-0.9) for *LRRK2 vs SNCA*, and 0.82 (95% CI: 0.81-0.82) for *PRKN* vs *SNCA* (Figure 4A). Per-class correct assignment was 65% for *PRKN*, 82% for *LRRK2*, and 60% for *SNCA* (Figure 4B). Misclassifications occurred most frequently between *PRKN* and *SNCA*, whereas *LRRK2* showed the highest correct classification rate. Weighted SHAP values in the multi-class setting showed contributions of 83% mitochondria and 17% lysosomes for *PRKN*, 50% mitochondrial and 50% lysosomal for *LRRK2*, and 82% mitochondrial and 18% lysosomal for *SNCA* (Figure 4C-E).

**Figure 4.**
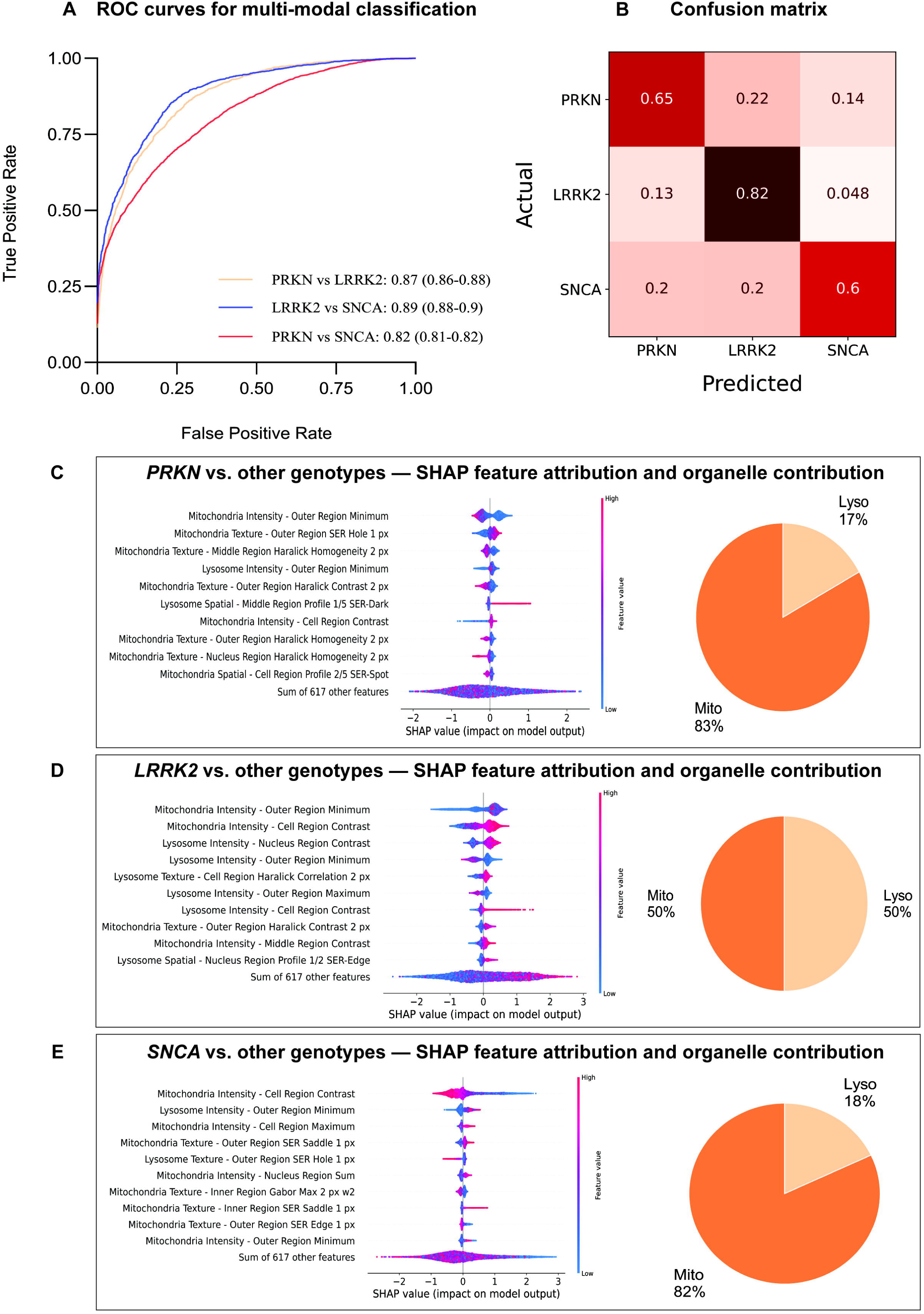
Multi-modal classification and organelle feature attribution across Parkinson’s disease genotypes. **(A)** ROC curves showing machine-learning based multi-class model performance in distinguishing *PRKN*, *LRRK2*, and *SNCA* neurons in the test dataset. Model separability was highest for *LRRK2* vs *SNCA* (AUC = 0.89, 95% CI 0.88–0.90), followed by *PRKN* vs *LRRK2* (AUC = 0.87, 95% CI 0.86–0.88) and *PRKN* vs *SNCA* (AUC = 0.82, 95% CI 0.81–0.82). **(B)** Confusion matrix summarising prediction accuracy across all genotypes. **(C–E)** SHAP feature attribution plots and organelle contribution charts for *PRKN* (C), *LRRK2* (D), and *SNCA* (E) neurons compared to other genotypes. Mitochondrial features dominated classification performance for *PRKN* and *SNCA* contributing 83% and 82%of explanatory power for *PRKN* and *SNCA* respectively, while lysosomal features were equal with mitochondrial features for *LRRK2*’s weighted SHAP explanations.

When compared with the binomial genotype-vs-control models (Figure 3), only small to moderate shifts (1–9%) were observed in organelle contributions between the multi-class and binomial models. *PRKN* showed a slight increase in mitochondrial contribution (82% to 83%) with a corresponding decrease in lysosomal contribution (18% to 17%). *LRRK2* showed a moderate decrease in mitochondrial contribution (59% to 50%) with a corresponding increase in lysosomal contribution (41% to 50%). *SNCA* showed a moderate increase in mitochondrial contribution (76% to 82%) and a corresponding decrease in lysosomal contribution (24% to 18%). Across both binomial and multi-class models, mitochondrial features accounted for the largest relative share of SHAP-based feature attribution for *PRKN* and *SNCA*, while *LRRK2* showed balanced contribution for multiclass classification.

### Robustness of morphomics across laboratories

To test the robustness of the assay, we compared morphomic predictions generated from two independent laboratories using different dyes and imaging conditions. Lab 1 used Mitotracker Red with 40× magnification, while Lab 2 used TMRM with 20× magnification. Despite these methodological differences, classifier performance was similar across the two sites.

In Lab 1, the classifier achieved an overall test accuracy of 66% with an average pairwise AUC of 0.83 (Figure 5A). Pairwise AUCs were strong for *PRKN* vs *LRRK2* (0.84, 95% CI: 0.82-0.85) and *LRRK2* vs *SNCA* (0.86, 95% CI: 0.85-0.87), while *PRKN* vs *SNCA* was more challenging (0.79, 95% CI: 0.78-0.79). Correct classification rates were highest for *LRRK2* (77%), followed by *PRKN* (63%), with *SNCA* showing greater misclassification (57% correctly classified).

**Figure 5.**
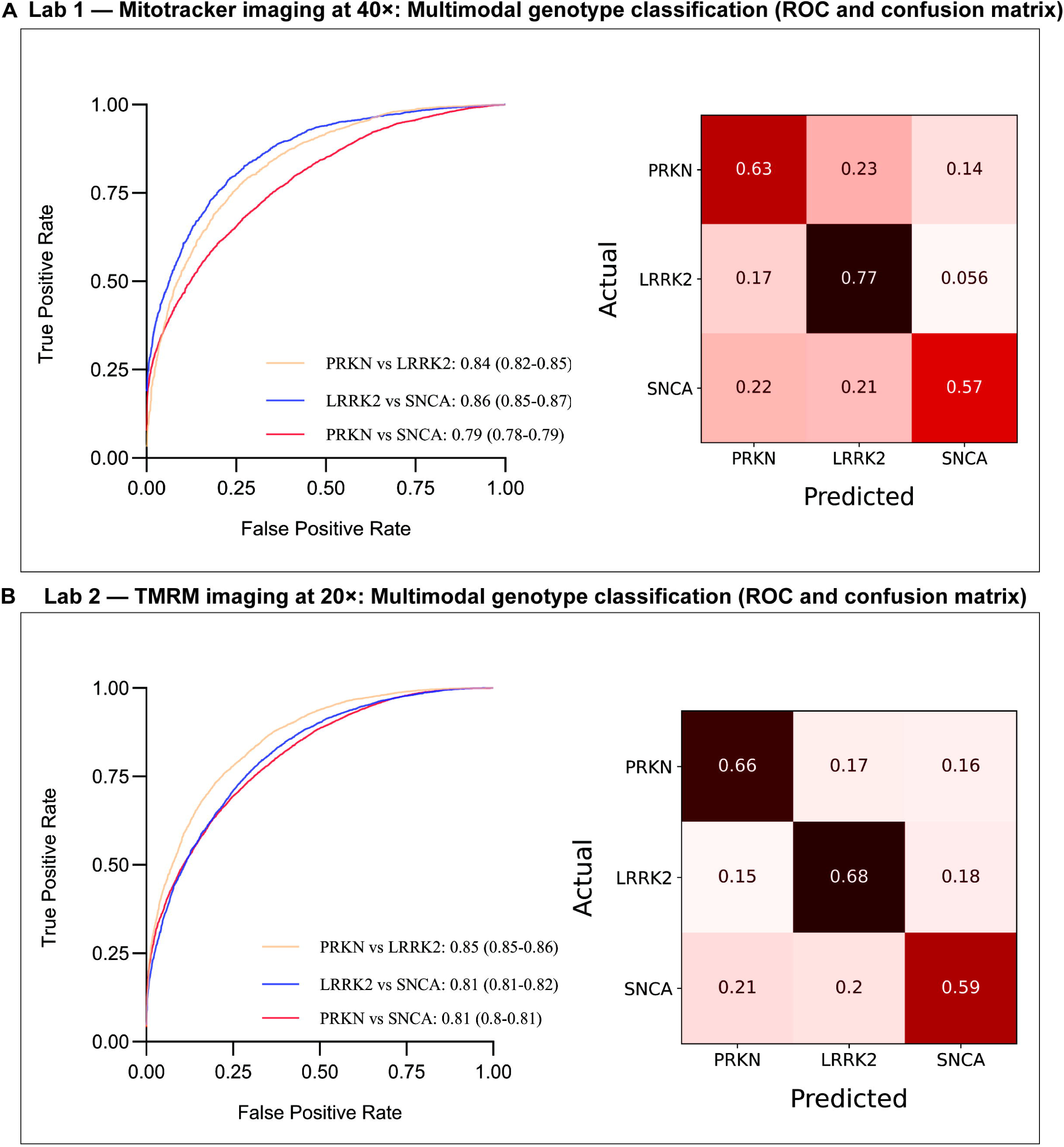
Cross-laboratory validation of multimodal genotype classification in Parkinson’s disease neurons. **(A)** Lab 1: Mitotracker imaging at 40× magnification. ROC curves and confusion matrix showing multi-class classification performance for *PRKN*, *LRRK2*, and *SNCA* neurons in the test dataset. Model separability was highest for *LRRK2* vs *SNCA* (AUC = 0.86, 95% CI 0.85–0.87), followed by *PRKN* vs *LRRK2* (AUC = 0.84, 95% CI 0.82–0.85) and *PRKN* vs *SNCA* (AUC = 0.79, 95% CI 0.78–0.79). **(B)** Lab 2: TMRM imaging at 20× magnification. ROC curves and confusion matrix showing consistent classification trends in the test dataset across imaging modalities, with comparable performance for *PRKN* vs *LRRK2* (AUC = 0.85, 95% CI 0.85–0.86), *LRRK2* vs *SNCA* (AUC = 0.81, 95% CI 0.81–0.82), and *PRKN* vs *SNCA* (AUC = 0.81, 95% CI 0.80–0.81).

In Lab 2, similar results were obtained (Figure 5B). The classifier reached an overall accuracy of 64%, with pairwise AUCs of 0.85 (95% CI: 0.85-0.86) for *PRKN* vs *LRRK2*, 0.81 (95% CI: 0.81-0.82) for *SNCA* vs *LRRK2*, and 0.81 (95% CI: 0.81-0.82) for *PRKN* vs *SNCA*. Correct classification was again strongest for *LRRK2* (68%) and *PRKN* (66%), while *SNCA* remained the least distinct (59%).

Together, these findings demonstrate that the morphomics workflow is robust across laboratories, remaining stable despite differences in staining dyes and imaging magnification. The reproducibility of predictions supports the generalisability of the assay and strengthens confidence in its biological readouts.

## Discussion

This study establishes mitochondrial and lysosomal morphomics as a robust framework for distinguishing genetic PD patient cells from controls and for resolving genotype-specific differences. By combining high-content imaging, machine learning, and SHAP-based attribution, we demonstrate that disease- and genotype-specific profiles can be consistently detected at single-cell resolution, and that these signals can be attributed directly to mitochondria and lysosomes.

At the disease level, PD-derived neurons were reliably separated from controls, confirming that morphomic features capture convergent cellular alterations shared across patients. Both mitochondrial and lysosomal features contributed to classification, however, mitochondrial features consistently accounted for a greater proportion of the model’s explanatory signal based on SHAP attribution. These changes are expected as in PD there is increase mitochondrial fission and reduced mitochondrial function and *LRRK2* mutations are known to increase lysosome tubulation and swelling [2]. The mitochondrial dyes used identify mitochondrial membranes and membrane potential, and the lysosomal dye used identifies lysosomal morphology and acidification, so the changes identified in the present study are consistent with known disease biology but indicate at a single-cell level that these abnormalities are widespread across neurons. At the genotype level, organelle-specific biases were apparent but broadly aligned with known biology: As shown by SHAP analysis in figure 3 *PRKN* cells were defined most strongly by mitochondrial descriptors, *LRRK2* cells showed the greatest relative lysosomal involvement, and *SNCA* cells displayed a more intermediate profile. These organelle-level “fingerprints” enabled robust discrimination between genotypes, while indicating a shared mitochondrial-dominant morphomic profile in *PRKN* and *SNCA* relative to other genotypes (Figure 4). Extending beyond binary classification, multi-class modelling showed that genotypes can also be distinguished directly from one another, albeit with lower accuracy than in genotype-vs-control comparisons. Importantly, organelle-level contributions were largely stable across modelling frameworks, with only small (1-9%) shifts in mitochondrial versus lysosomal weighting.

In line with these morphomic signatures, our earlier functional analyses of the same cortical neuron lines demonstrated that all three genotypes exhibited significantly reduced mitochondrial respiration, whereas lysosomal glucocerebrosidase activity was most severely impaired in *LRRK2* neurons [23]. The dominance of mitochondrial descriptors across prediction models therefore reflects a convergent mitochondrial deficit that was captured independently by oxygen consumption rate assays. Conversely, the enhanced contribution of lysosomal features to *LRRK2* classification aligns with its pronounced lysosomal dysfunction and highlights a distinct morphomic profile in this genotype. Notably, in the present morphomics analysis, mitochondrial features contributed most strongly to *PRKN* classification, a pattern that was not evident in the bulk functional assays where all genotypes showed comparable reductions in respiration. This suggests that the single-cell resolution of the morphomics approach is more sensitive to detecting genotype-specific mitochondrial alterations than population-level assays such as oxygen consumption rate measurements[24]. This heightened sensitivity aligns with the established role of *PRKN* in maintaining mitochondrial quality control, making subtle mitochondrial defects in *PRKN* neurons more readily detectable through morphomic profiling.

Our results complement and extend prior iPS cell-based morphomic studies, where both mitochondria and lysosomes have been implicated as key contributors to PD phenotypes. Using profiled iPS cell-derived cortical neurons carrying *SNCA* triplication and *PINK1* mutations, alongside chemically induced PD-like models including rotenone (complex I dysfunction), oligomycin/antimycin (induced mitophagy), and α-synuclein oligomers (proteinopathy), their classifier achieved an overall accuracy of 82% for a PD phenotype[20]. SHAP analysis revealed that mitochondrial features, particularly TMRM-based texture descriptors, dominated prediction across most subtypes, whereas lysosomal features were most prominent in *SNCA* triplication[20]. Similarly, Vuidel *et al* reported that *LRRK2* G2019S dopaminergic neurons display elevated α-synuclein levels, reduced dendritic complexity, abnormal mitochondrial morphology (reduced membrane potential and increased compactness), and altered lysosomal features. Using machine-learning classifiers, these cells were separated from isogenic and unrelated controls, and drug interventions (*LRRK2* inhibitors, PKC agonists) shifted morphologies towards controls. While these studies established proof-of-concept that morphomics can detect genotype-specific phenotypes, they relied on only one or two patient lines per genotype, limiting their generalisability. In contrast, our analysis of 13 iPS cell-derived neuronal lines across *PRKN*, *LRRK2*, *SNCA*, and matched controls demonstrates that organelle-specific signatures are robust across diverse donors and more sensitive than bulk functional assays[23]. Moreover, by validating morphomic results in two independent laboratories using different mitochondrial dyes and imaging conditions, we show that morphomic predictions are stable despite methodological variation. Together, these advances move morphomics beyond proof-of-concept to a reproducible and scalable framework.

Morphomic approaches have also been applied beyond neurons. Schiff *et al*. (2022) profiled fibroblasts from 46 PD patients (sporadic, *LRRK2*, *GBA*) and 45 controls using a Cell Painting assay (>3,000 features across multiple organelle and cellular compartments)[25]. PD fibroblasts were distinguished from controls with AUCs of 0.76-0.89, and donor-specific morphologies remained stable across biopsies collected 3-6 years apart, indicating persistent, reproducible signatures. Anthony *et al*. (2020) analysed fibroblasts from 41 idiopathic PD patients and 21 controls and found increased mitochondrial fragmentation and reduced branching in most parameters; classification improved under FCCP-induced stress (AUC 0.87 vs 0.83), showing how stress can unmask latent defects[26]. These fibroblast studies demonstrate the broad applicability of morphomics and the stability of donor-specific profiles. However, fibroblasts provide only an indirect view of PD. By focusing on iPS cell-derived neurons, our study interrogates disease-relevant cell types and links genotype-specific differences directly to mitochondrial and lysosomal alterations at single-cell resolution.

Notably, while each genotype displayed distinct organelle-level biases, the presence of both mitochondrial and lysosomal alterations across all groups reinforces the concept of shared pathogenic pathways in PD. As discussed in the introduction and supported by our findings, these genotypes converge on similar organellar dysfunctions but differ in the relative extent to which each pathway is affected. Identifying the dominant contributor within each genotype, whether mitochondrial or lysosomal, may therefore provide a rational basis for selecting and testing pathway-specific therapeutic interventions. For example, compounds enhancing mitophagy or mitochondrial biogenesis may be particularly effective in *PRKN*-related disease, whereas strategies targeting lysosomal function or autophagic flux may hold greater promise for *LRRK2*- or *SNCA*-related disease. Combination approaches targeting both pathways may further enhance therapeutic efficacy. Moreover, this morphomics-based framework could offer a path to identifying cellular pathology in idiopathic patients lacking defined genetic information, thereby extending cell pathology targeted strategies to the majority of PD cases. This framework could guide drug screening and precision therapeutic design by aligning molecular pathology with targeted rescue mechanisms.

Our work has certain limitations. Genotype overlap, particularly in mitochondrial morphomics between *PRKN* and *SNCA*, highlights persistent challenges in resolving PD heterogeneity. Incorporating additional cellular compartments, markers such as total and phosphorylated α-synuclein, or stress paradigms may enhance discriminatory power. While mitochondria consistently dominated contributions, lysosomes provided complementary and non-redundant information, especially in *LRRK2* and *SNCA*, underscoring the importance of multi-organelle profiling.

In summary, this study advances morphomics from proof-of-concept to a reproducible and generalisable platform for PD research in iPS cell-derived neurons. By leveraging a large multi-donor cohort, cross-laboratory validation, and organelle-level feature attribution, we establish a scalable framework with potential applications in patient stratification, biomarker discovery, and drug development.

## Methods

### Cell lines

All 13 iPS cell lines described in this study, *LRRK2* R1441G (n = 3), *SNCA* A53T (n = 3), *PRKN* lof mutations (n = 3), and neurologically normal controls (n = 4) have been described in detail in our recent publication [23]. Experiments on human iPS cells were approved by the Human Research Ethics Committee at the University of Sydney (2017/094) and University of Melbourne (2020-1956049 & 2025-33900-73031-3).

### Cortical neuron differentiation

Cortical neural progenitors were generated from iPS cells according to the methods of [27]. At day 25 post differentiation, vials of cortical progenitors were cryopreserved and retained in vapour phase liquid nitrogen until required. Neural progenitor cells were then thawed in the presence of 10 µM Y27632 (Stem Cell Technologies, Cat#72304) and plated into PhenoPlate 96 well plates (Perkin Elmer) pre-coated with 15 µg/ml Poly-L-ornithine (Sigma, Cat# P4957) and 10 µg/ml Mouse Laminin (Sigma, Cat# L2020) at a density of 10,000 cells per well. Cells were initially cultured in cortical base media comprised of a 1:1 mix of Dulbecco’s Modified Eagle Medium F12 (DMEM/F12) and Neurobasal media containing 0.5x B27 supplement, 0.5x N2 supplement, 0.5x ITS-A, 0.5x non-essential amino acids, 0.5x glutamax, 50 U/ml penicilin/streptomycin and 49.5 µM β-mercaptoethanol (all from Thermofisher). The Y27632 was removed from culture media at day1 and replaced by the cortical base media only. From day3, the media was changed to cortical maturation media comprised of a 1:1 mix of DMEM/F12 and Neurobasal media containing 1x B27 supplement, 1x N2 supplement, 1x ITS-A, 1x non-essential amino acids, 0.5x glutamax and 50 U/ml penicilin/streptomycin (all from Thermofisher) and further supplemented with 40 ng/ml BDNF, 40 ng/ml GDNF (both from Stem Cell Technologies), 50 µM Dibutyryl cAMP (Stem Cell Technologies), 200 nM ascorbic acid (Sigma), 100 ng/ml Mouse Laminin (Sigma) and 10 µM of the γ-secretase inhibitor DAPT (Sigma). Cells were differentiated for 12 days in cortical maturation media, with 90% of the spent media replaced every second day with fresh media. At day 15 post-thaw, a total of 40 days of differentiation, the cells were used for experiments.

### Immunocytochemistry for charecterisation

Differentiated cortical cells from all lines were stained with COUP-TF-interacting protein 2 (CTIP2) to identify layer V cells, T-box brain protein 1 (TBR1) to identify deep-layer V and VI cells together with microtubule-associated protein 2 (MAP2) as a general neuronal marker. Cells were stained with antibodies against MAP2, TBR1 and CTIP2 to identify mature cortical neurons. Immunostaining was performed using the CytoFix/CytoPerm fixation and permeabilization kit (BD Biosciences) as previously described (PMID 34404843). Briefly, cells were fixed in Cytofix for 20 min, washed three times (5 min each), and permeabilised in Cytoperm buffer for 30 min. Primary antibodies (Supplementary Table 1) were applied for 1 h, followed by three washes (5 min each) and incubation with secondary antibodies (Supplementary Table 2) for 30 min. Hoechst nuclear dye (ThermoFisher) was included during the final 10 min of secondary antibody incubation. After three additional washes, plates were sealed with parafilm and stored at 4 °C in the dark until imaging. One-way ANOVA comparing each genotype to control did not reveal any statistically significant differences in the expression of TBR1 and CTIP2 markers (Supplementary figure 2).

### Live-cell assays

Live-cell imaging assays were performed to assess mitochondrial and lysosomal phenotypes. 96 well-plates were imaged on the Opera Phenix high-content imager (PerkinElmer). Working dilutions of live-cell reagents were prepared in culture media and added to wells after removal of old media. Details of all reagents used are mentioned in Supplementary Table 3

For the mitochondrial and lysosomal assays described in Figures 1-4, cells were incubated with Mitotracker Deep Red (Thermo Fisher, M22426), lysosomal staining reagent Cytopainter (Abcam, ab176827) and Cytopainter Green (abcam, 138891) dyes for 60 min. Staining with Hoechst 33342 (Thermo Scientific, 62249) to label nuclei was then performed for 10 min and cells washed with media and imaged immediately.

For mitochondrial TMRM phenotyping described in Figure 4, cells were labelled with Calcein (Invitrogen, C3100MP) to identify viable cells, TMRM (Invitrogen, T-668) to assess mitochondrial membrane potential, and Hoechst 33342 (Thermo Scientific, 62249). Cells were incubated with probes for 30 min, washed twice with media, and imaged immediately.

### High content imaging

Images were acquired on the Opera Phenix PLUS high-content screening system (Revvity) using either 20× or 40× water-immersion Zeiss Plan Apochromat objectives (20×, NA 1.0, WD 1.7 mm; 40×, NA 1.1, WD 0.62 mm). Detection was performed using a monochrome sCMOS camera (2160 × 2160 pixels, 6.5 μm pixel size, quantum efficiency >80%, read noise <1 e⁻, dynamic range ∼33,000:1). The system was equipped with a microlens-enhanced dual-layer spinning disk Nipkow confocal device (50 μm pinhole size, 555 μm inter-pinhole distance), dual excitation paths to minimise cross-talk, and laser-based illumination. All images were acquired in 16-bit TIFF format.

Image acquisition and downstream analysis were performed using Harmony software (v5.2, Revvity). For each experiment, details including objective magnification, average number of cells imaged, fluorescence channels, number of fields of view, Z-step size, and number of planes per Z-stack are provided in Supplementary Table 4. For live-cell imaging assays, the number of Z-steps per stack was minimised to enable completion of each experiment within 30 minutes, ensuring consistency across experimental conditions.

### Image data extraction

Image segmentation and feature extraction were performed using Harmony High-Content Analysis software version 5.2 (Revvity) following a Cell Painting style workflow[21]. Nuclei were segmented from the Hoechst channel, and cytoplasmic boundaries were defined using Calcein (for live-cell assays) or Cytopainter (for fixed-cell assays). To reduce segmentation errors from overlapping neuronal processes, cytoplasmic masks were restricted to include only the soma and proximal neurites.

Feature extraction was performed on the mitochondrial (Mitotracker or TMRM) and lysosomal (Lysotracker Deep Red) channels using the “Calculate Cell Painting Properties” module with a standard SER scale of 1. Each organelle contributed 660 single-cell features, grouped into four main categories: intensity (mean, SD, quantiles, contrast), morphology (area, shape, length, perimeter), texture (SER, Haralick, and Gabor descriptors capturing spot, edge, and granularity patterns), and spatial (radial distribution, symmetry, compactness, and perinuclear enrichment).

Features were quantified across five concentric cellular regions - nuclear, inner, middle, cell, and outer capturing the radial organisation of organelles. Border-touching objects were excluded, and the resulting datasets were exported as tab-delimited text files with embedded metadata (cell line, plate, and batch) for downstream machine-learning analysis.

### Data analysis

Highly correlated features (Spearman’s ρ > 0.9) were removed, reducing the dataset from 1320 features to 627 (325 Lysosomal, 302 Mitochondrial). Cells with missing feature values and wells with cell counts below 200 were excluded from analysis. In total 9% and 18% of cells were removed from laboratory 1 and 2 respectively due to missing values and low well cell counts

The dataset was then split into training (60%), validation (20%) and test (20%) datasets. To account for batch effects features were scaled to training set controls in each batch separately using a standard scaler. To maintain class balance for training the classification models, the training dataset was randomly down sampled to the minority class for each prediction model (Supplementary Table 5). ML predictions were made using a Light Gradient-Boosting Machine (LGBM) classifier with Python using the LightGBM package. Classifiers were trained in the training datasets and n_estimators was optimised in the validation set by enabling 20 rounds of early stopping, using the balanced log loss in the validation sets as the evaluation metric. All other LGBM hyperparameters used default values. The final results are the predictions in the test datasets.

Graphs were generated in GraphPad Prism and Python. Multiclass receiver operator curves were generated using the interpolated average of the receiver operator curves for a given pairwise comparison. AUC-ROC confidence intervals were calculated using bootstrapping resampling.

### SHAP-weighted organelle attribution

To determine the relative contribution of mitochondria and lysosomes to classification, we analysed SHAP (SHapley Additive exPlanations) values from each trained model. SHAP values were calculated on class balanced datasets. For each comparison (e.g., controls vs. PD, controls vs. PRKN), we extracted the absolute mean SHAP value for all features. Features were ranked by SHAP values, and the top 10 features were used for organelle-level attribution.

The weighted contribution of each organelle was calculated as the sum of SHAP values for features belonging to that organelle, divided by the total SHAP values across the top *10* features. Mitochondrial and lysosomal contributions therefore represent the proportion of explanatory power attributable to features from each compartment. This approach accounts for both the number of features and their weighted influence on the classifier’s decision, in contrast to simple feature counts.

### Data availability

The data, protocols, and key lab materials used and generated in this study are listed in a Key Resources Table alongside their persistent identifiers. All data and analysis code generated in this study and the Key Resources Table are publicly available via Zenodo repository: 10.5281/zenodo.18323330.

The Zenodo repository includes single-cell feature datasets derived from high-content imaging, provided as spreadsheet files. These include raw feature tables from individual experimental batches and laboratories, as well as batch-normalised and combined datasets used for downstream analysis. Data preprocessing, statistical analysis, machine-learning classification, and SHAP-based feature attribution were performed using Jupyter Notebook scripts deposited in the same Zenodo record. A README file is included to describe the contents of the repository, file structure, and the purpose of each dataset and analysis notebook. A feature name reference file is also provided to map original feature names generated by the imaging software to the harmonised feature names used in this study. Raw microscopy image files are stored separately due to their size and are available from the corresponding author upon reasonable request.

## Acknowledgements

This work was supported in part by *Aligning Science Across Parkinson’s* (ASAP-000497) through the Michael J. Fox Foundation for Parkinson’s Research (MJFF). CLP and GMH are NHMRC Leadership Fellows. To facilitate open access, the authors have applied a CC BY public copyright license to all Author Accepted Manuscripts arising from this submission. Data used in the preparation of this article were obtained from the Parkinson’s Progression Markers Initiative (PPMI) database (www.ppmi-info.org/access-data-specimens/download-data). PPMI is a public-private partnership funded by The Michael J. Fox Foundation for Parkinson’s Research and its industry partners (see the full list at https://www.ppmi-info.org/about-ppmi/who-we-are/study-sponsors) — provides open-access clinical and imaging data to advance Parkinson’s research. The authors also acknowledge the Sydney Cytometry Core Research Facility, a joint initiative of the Centenary Institute and the University of Sydney, for access to the Opera Phenix high-content imaging system.

## Author contributions

Conceived the idea and secured funding – D.K., G.M.H., C.S., L.T., C.P., J.J. Planned the experiments – G.W., N.D., C.S. Generation and Q.C. of iPS cell lines – D.O., R.D., D.S., C.S., C.P. Performed the experimental work – Y.L., J.C., G.W., A.L.G., M.Z., D.A.B., C.P., T.F. Data analysis and figures – G.W., M.P., R.S., N.D., J.C. Drafted the manuscript –G.W., M.P., Y.L., N.D., C.S., G.M.H. Edited and approved the final version – all authors.

---

**Supplementary figure 1.**
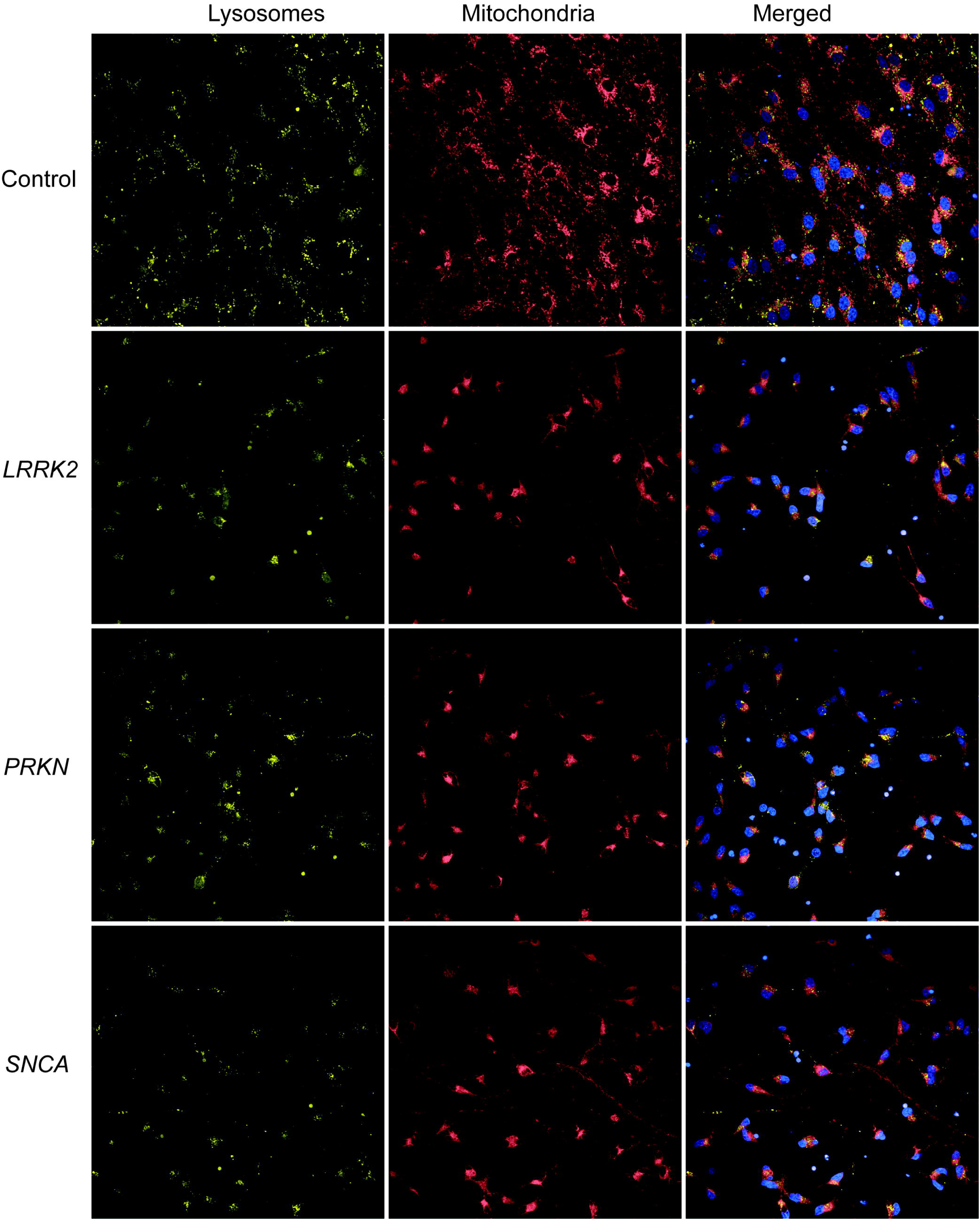
Representative mitochondrial and lysosomal images across control and Parkinson’s disease genotypes. Representative fluorescence images of iPS-derived neurons from control, *LRRK2*, *PRKN*, and *SNCA* groups labelled with MitoTracker (red) to visualise mitochondria and LysoTracker (yellow) to visualise lysosomes, with merged images including nuclear staining (blue).

**Supplementary figure 2.**
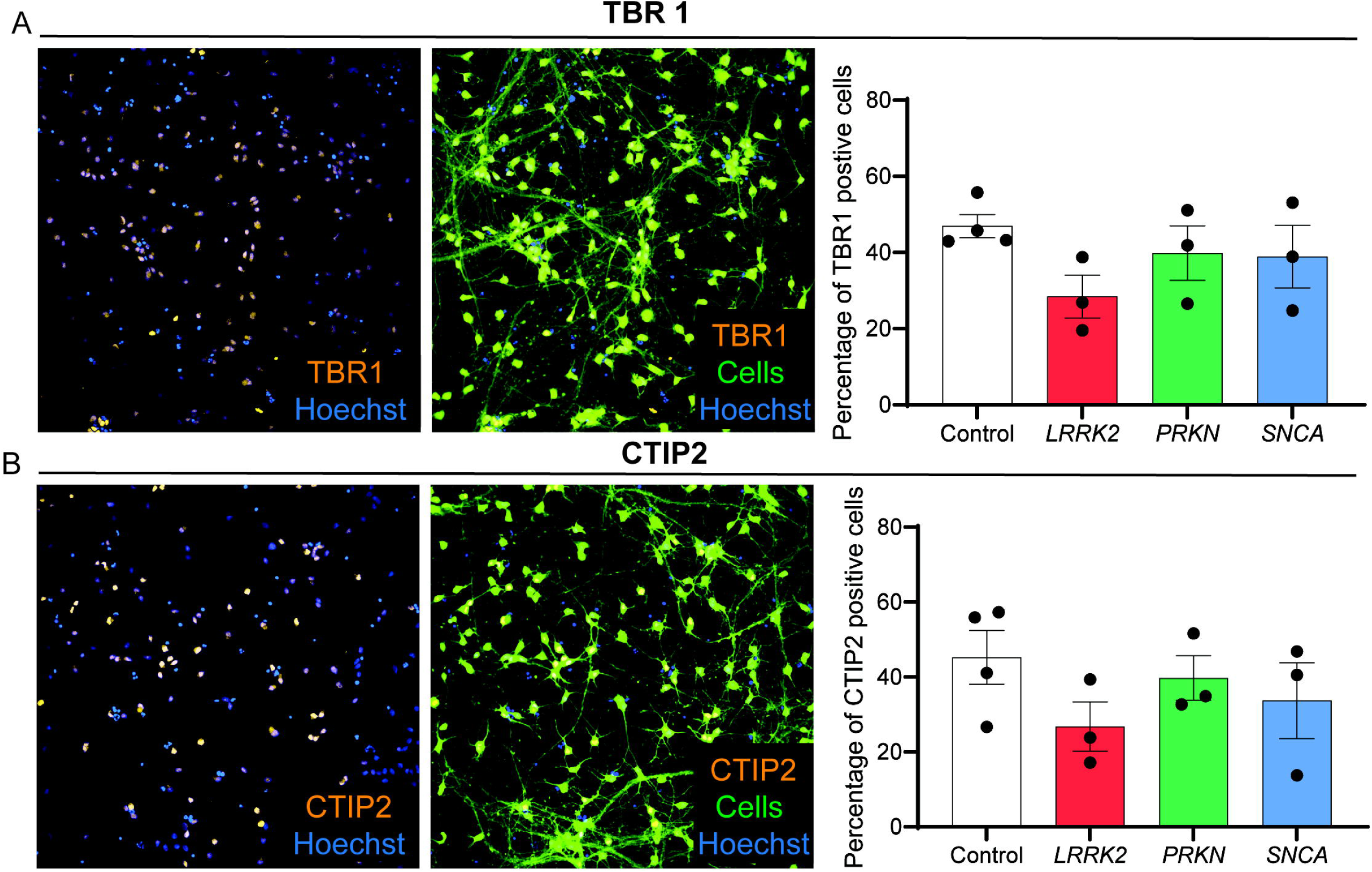
Cortical neuron characterization across control and Parkinson’s disease genotypes. Representative immunofluorescence images showing expression of (A) the layer VI cortical marker (A) TBR1 (yellow) and (B) layer V cortical marker CTIP2 (yellow) with Hoechst nuclear staining (blue) and the neuronal marker MAP2 (green), alongside quantification of the percentage of (A) TBR1 and (B) CTIP2 positive cells in control, *LRRK2*, *PRKN*, and *SNCA* iPSC-derived neurons. Data are shown as mean ± SEM.

