## Supplementary tables 1-5 for "Machine learning–based image analysis of Parkinson’s disease iPS cell-derived neurons predicts genotype and reveals mitochondria–lysosome abnormalities"

| **Antibody** | **Company** | **Catalogue no** | **Species** | **Dilution** |
| --- | --- | --- | --- | --- |
| TBR1 | Abcam | ab31940 | rabbit | 1:1000 |
| CTIP2 | Abcam | ab18465 | rat | 1:1000 |
| MAP2 | Abcam | ab254143 | mouse | 1:1000 |

**Supplementary Table 1. Antibodies. Details for primary antibodies employed for immunocytochemistry.**

| **Antibody** | **Company** | **Catalogue no** | **Species** | **Dilution** |
| --- | --- | --- | --- | --- |
| Goat anti-mouse-488 | Invitrogen | A-11029 | Goat | 1:500 |
| Goat anti-rabbit-488 | Invitrogen | A-11008 | Goat | 1:500 |
| Goat anti-rabbit-594 | Invitrogen | A-11012 | Goat | 1:500 |
| Goat anti-rat-647 | Invitrogen | A-48272 | Goat | 1:500 |
| Goat anti-rat-488 | Invitrogen | A-11006 | Goat | 1:500 |

**Supplementary Table 2. Secondary Antibodies. Details for secondary antibodies employed for immunocytochemistry.**

| **Serial no** | **Assay** | **Stain / reagent** | **Company** | **Catalogue no** | **Final concentration** |
| --- | --- | --- | --- | --- | --- |
| 1 | Mitochondrial TMRM assay (Figure 4) | Tetramethylrhodamine, Methyl Ester, Perchlorate (TMRM) | Invitrogen | T668 | 25 nM |
| 2 |  | Calcein | Invitrogen | C3100 | 500 nM |
| 3 |  | Hoechst | Thermo Scientific | 62249 | 1 µg/ml |
| 4 | Mitotracker and Lysotracker assay (Figures 1-4) | Mitotracker Deep Red | Thermo Fisher | M22426 | 100 nM |
| 5 |  | Cytopainter green cytoplasm marker | Abcam | ab138891 | 1x |
| 6 |  | Lysosomal staining reagent – orange- cytopainter | Abcam | ab176827 | 1x |
| 7 |  | Pureblu Hoechst 33342 | Bio-Rad | 1351304 | 2 µM |

**Supplementary Table 3**. Live cell imaging dyes used for mitochondria and lysosome live cell assay.

| **Assay** | **Lab 1 or 2** | **Number of batches (plates)** | **Objective** | **Number of wells per plate per cell line** | **Z-stack: Number of planes** | **Z-stack: Step size** | **Number of fields of view imaged per well** | **Channel number (pre-defined filter)** | **Imaged** |
| --- | --- | --- | --- | --- | --- | --- | --- | --- | --- |
| Mitochondria and lysosome assay | Lab 1 | 3 | 40x | 3 | 3 | 0.5µm | 25 | Ch1 (Lysotracker Red DND) | Lysosomes |
|  |  |  |  |  |  |  |  | Ch2 (Mitoctracker DeepRed) | Mitochondria |
|  |  |  |  |  |  |  |  | Ch3 (Cytopainter) | Cytoplasm |
|  |  |  |  |  |  |  |  | Ch4 (Hoechst) | Nuclei |
|  |  |  | 40x | 3 | 3 | 0.5µm | 25 | Ch1 (Mitotracker Orange) | Lysosomes |
|  |  |  |  |  |  |  |  | Ch2 (Mitoctracker DeepRed) | Mitochondria |
|  |  |  |  |  |  |  |  | Ch3 (Cytopainter) | Cytoplasm |
|  |  |  |  |  |  |  |  | Ch4 (Hoechst) | Nuclei |
|  |  |  | 40x | 3 | 4 | 0.5µm | 25 | Ch1 (Lysotracker Orange DND) | Lysosomes |
|  |  |  |  |  |  |  |  | Ch2 (Mitoctracker DeepRed) | Mitochondria |
|  |  |  |  |  |  |  |  | Ch3 (Cytopainter) | Cytoplasm |
|  |  |  |  |  |  |  |  | Ch4 (Hoechst) | Nuclei |
| Mitochondria (TMRM) assay | Lab 2 | 3 | 20x | 3 | 3 | 2µm | 15 | Ch1 (Alexa 488) | Cytoplasm |
|  |  |  |  |  |  |  |  | Ch2 (Alexa 568) | Mitochondria (TMRM) |
|  |  |  |  |  |  |  |  | Ch3 (CellMask DeepRed) | Mitochondria (Mitotracker) |
|  |  |  |  |  |  |  |  | Ch4 (Hoechst) 33342 | Nuclei |
|  |  |  | 20x | 3 | 3 | 2µm | 15 | Ch1 (Alexa 488) | Cytoplasm |
|  |  |  |  |  |  |  |  | Ch2 (Alexa 568) | Mitochondria (TMRM) |
|  |  |  |  |  |  |  |  | Ch3 (Brightfield) | Brightfield |
|  |  |  |  |  |  |  |  | Ch4 (Digital Phase Contrast) | Digital Phase Contrast |
|  |  |  |  |  |  |  |  | Ch5 (Hoechst 33342) | Nuclei |
|  |  |  | 20x | 3 | 3 | 2µm | 15 | Ch1 (Alexa 488) | Cytoplasm |
|  |  |  |  |  |  |  |  | Ch2 (Alexa 568) | Mitochondria (TMRM) |
|  |  |  |  |  |  |  |  | Ch3 (Brightfield) | Brightfield |
|  |  |  |  |  |  |  |  | Ch4 (Digital Phase Contrast) | Digital Phase Contrast |
|  |  |  |  |  |  |  |  | Ch5 (Hoechst 33342) | Nuclei |

**Supplementary Table4**. Imaging acquisition parameters for mitochondrial and lysosomal assays.

|  | Training | Validation | Test |
| --- | --- | --- | --- |
| **PD vs Controls** |  |  |  |
| PD | 12560 | 7802 | 7803 |
| Controls | 12560 | 4187 | 4187 |
| **PRKN vs Controls** |  |  |  |
| Controls | 10504 | 4187 | 4187 |
| *PRKN* | 10504 | 3501 | 3501 |
| ***LRRK2* vs Controls** |  |  |  |
| *LRRK2* | 2365 | 788 | 788 |
| Controls | 2365 | 4187 | 4187 |
| ***SNCA* vs Controls** |  |  |  |
| *SNCA* | 10539 | 3513 | 3514 |
| Controls | 10539 | 4187 | 4187 |
| ***PRKN* vs *LRRK2* vs *SNCA*** |  |  |  |
| *PRKN* | 2365 | 3501 | 3501 |
| *LRRK2* | 2365 | 788 | 788 |
| *SNCA* | 2365 | 3513 | 3514 |
| ***PRKN* vs *LRRK2* vs *SNCA* (Laboratory 2)** |  |  |  |
| *PRKN* | 7407 | 5972 | 5972 |
| *LRRK2* | 7407 | 2469 | 2469 |
| *SNCA* | 7407 | 7919 | 7920 |

**Supplementary Table 5**. Cell counts used for training, validation, and test datasets for each classification model.
